# Evolutionary dynamics with directed updating rules

**DOI:** 10.64898/2026.11.26.769373

**Authors:** Wei Lu, Joshua B. Plotkin, Anzhi Sheng

## Abstract

Evolutionary game theory describes how the frequencies of strategies change in a population when individuals imitate others who have higher payoffs. Models of imitation typically assume that updating is undirected, so that any individual may adopt the strategy of any other. But in many systems some replacements occur in one direction only: in a food chain, for example, a predator species can replace its prey but not the reverse. Here we study replicator dynamics with directed updating rules in a generalized Rock-Paper-Scissor game, in which each strategy can replace only one of the other two. Under undirected updating, increasing the payoff that Paper receives against Rock produces a population in which Scissors is close to fixation. Under directed updating, by contrast, the frequencies of all three strategies oscillate around an interior state. Moreover, when each strategy can replace only the strategy that beats it, increasing the payoff of Paper against Rock decreases the equilibrium frequency of Scissors, which is the opposite of its effect at the Nash equilibrium of the game. This reversal accompanies a reversal in the direction in which the population state rotates around the interior equilibrium. Our results indicate that the direction of strategy updating, and not only the payoff structure of the game, determines which strategies are common at equilibrium.

## 1 Introduction

Evolutionary game theory, which originated in the study of animal conflict (Maynard Smith & Price 1973), describes how the frequencies of behaviors, or strategies, change in a population as individuals interact and imitate one another (Helbing 1992; Hofbauer & Sigmund 2003; Sandholm 2010). In a common formulation of imitation, an individual compares its payoff with the payoff of a randomly chosen role model, and it adopts the strategy of the role model with a probability that increases with the payoff difference (Schlag 1998; Traulsen et al. 2006, 2007, 2010). In a large population and under weak selection, this pairwise comparison process yields the replicator equation (Traulsen et al. 2005), whose interior equilibria are Nash equilibria of the game (Taylor & Jonker 1978; Schuster & Sigmund 1983; Hofbauer & Sigmund 2003; Cressman & Tao 2014). And so, in this classical setting, the long-term frequencies of strategies are predicted from the payoff matrix alone.

The Rock-Paper-Scissor game is the simplest game with cyclic dominance: Rock beats Scissors, Scissors beats Paper, and Paper beats Rock (Hofbauer & Sigmund 2003; Szolnoki et al. 2014). Cyclic dominance of this kind occurs in natural populations, for example among strains of bacteria and among the male morphs of the side-blotched lizard (Sinervo & Lively 1996; Kerr et al. 2002; Kirkup & Riley 2004). In the replicator dynamics of a zero-sum Rock-Paper-Scissor game, the strategy frequencies oscillate around an interior equilibrium whose position is determined by the payoffs (Hofbauer & Sigmund 1998). At this equilibrium, increasing the payoff that one strategy receives against the strategy it beats increases the equilibrium frequency not of that strategy, but of the strategy that beats it. A related effect occurs in models and experiments of cyclic competition among species, in which an increase in the competitive ability of one species lowers its own abundance (Frean & Abraham 2001; Nahum et al. 2011).

But models of imitation dynamics typically assume that updating is undirected: any strategy can replace any other, provided the payoff difference favors it. In many systems, some replacements occur in one direction only. In a food chain, a predator can replace its prey, but the prey cannot replace the predator (Volterra 1926; Holling 1959). In cancer immunology, macrophages can engulf dead tumor cells, but the reverse does not occur (Asano et al. 2011; Baghdadi et al. 2013; Zhou et al. 2020). In an epidemic, individuals pass from susceptible to infected to recovered, but not in the reverse order (Kermack & McKendrick 1927; Anderson & May 1991; Hethcote 2000). And in social learning, behaviors are often copied from individuals of higher status or greater experience, and seldom in the reverse direction (Henrich & Gil-White 2001; Henrich & Broesch 2011; Chudek et al. 2012). In each of these systems, the set of possible replacements is restricted in direction, separately from any payoffs.

Restrictions on who can replace whom have been studied by placing individuals on a graph (Ohtsuki et al. 2006; Allen et al. 2017; Su et al. 2022; Sheng et al. 2026), including directed graphs on which one individual can replace another but not the reverse (Lieberman et al. 2005; Masuda 2009; Masuda & Ohtsuki 2009). In those models the restriction depends on the positions of individuals, and every strategy can in principle replace every other. How a restriction on which strategy can replace which changes the outcome of evolution in a well-mixed population is less well understood, leaving basic questions open. Does restricting the direction of replacement change whether one strategy approaches fixation? Does it change the direction in which the population state rotates around the interior equilibrium? And does it change how the equilibrium frequencies of strategies depend on the payoffs?

Here we study pairwise comparison updating with a replacement matrix that specifies which strategy can replace which, in a generalized Rock-Paper-Scissor game. We derive the replicator equations for an arbitrary replacement matrix, and we compare undirected updating with two directed rules: a clockwise rule, under which each strategy can replace only the strategy it beats in the game, and a counterclockwise rule, under which each strategy can replace only the strategy that beats it. Under undirected updating, increasing the payoff of Paper against Rock produces a population in which Scissors is close to fixation. Under both directed rules, by contrast, we find that the frequencies of all three strategies oscillate around an interior state and remain at intermediate values. Under counterclockwise updating, the population state rotates in the order opposite to the cycle of dominance in the game, and a larger payoff of Paper against Rock lowers the equilibrium frequency of Scissors, which is the opposite of its effect at the Nash equilibrium. We examine the rates at which each strategy replaces each other strategy at a reference state, to explain how the equilibrium frequency of Scissors depends on the payoff of Paper against Rock, and we find that the reversal under counterclockwise updating also occurs under strong selection. Our results indicate that the direction of strategy updating, and not only the payoff structure of the game, determines which strategies are common at equilibrium.

## 2 Model

We consider a large, well-mixed population in which each individual plays a generalized Rock-Paper-Scissor game with every other individual (Fig. 1a). Each individual uses one of three strategies: Rock (R), Paper (P), or Scissors (S). Paper beats Rock, Rock beats Scissors, and Scissors beats Paper. When Paper meets Rock, the Paper player gains *b* and the Rock player loses *b*; when Rock meets Scissors, the Rock player gains *a* and the Scissors player loses *a*; and when Scissors meets Paper, the Scissors player gains 1 and the Paper player loses 1. Two players who use the same strategy each receive 0. The game is therefore zero-sum: in every interaction, one player’s gain equals the other player’s loss. The parameters *a >* 0 and *b >* 0 are the payoffs of Rock against Scissors and of Paper against Rock, measured relative to the payoff of Scissors against Paper.

**Figure 1:**
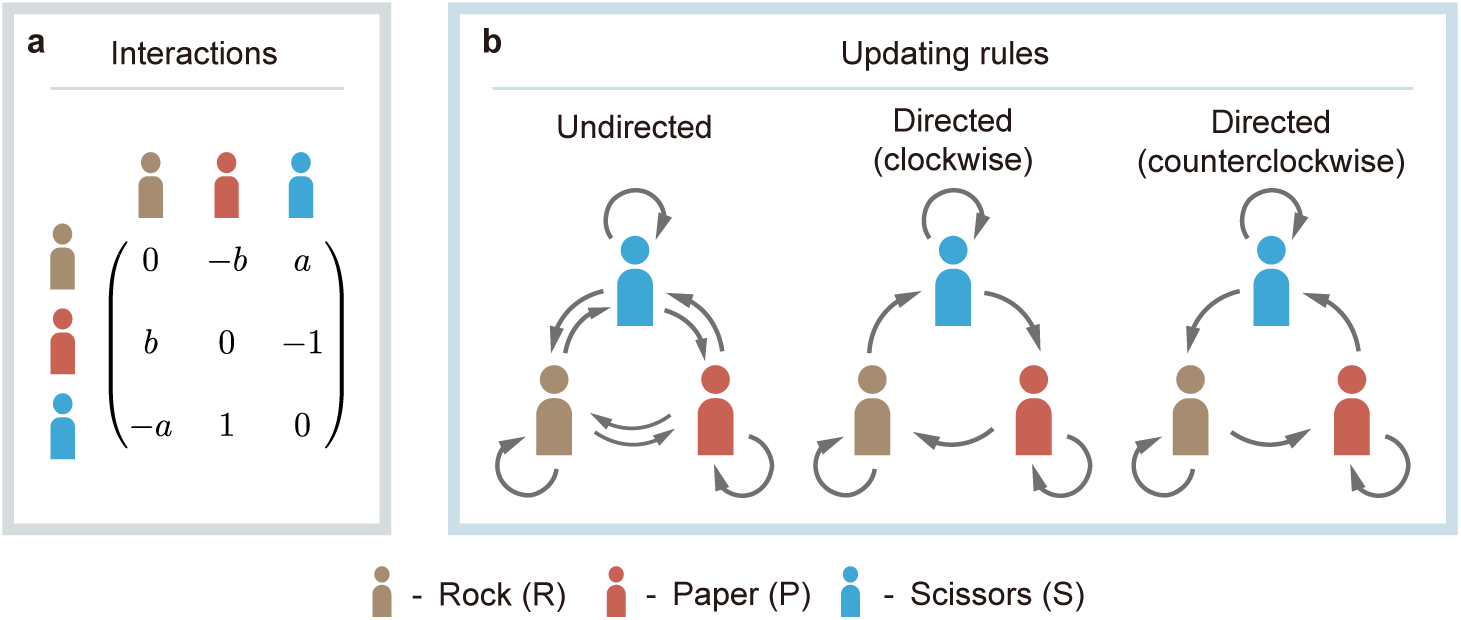
A generalized Rock-Paper-Scissor game with undirected and directed updating. We consider a large, well-mixed population in which individuals play a generalized Rock-Paper-Scissor game. **a**, Payoff matrix of the game among Rock (R, brown), Paper (P, red), and Scissors (S, blue). Each entry is the payoff to the row strategy against the column strategy. **b**, Three updating rules. An arrow from one strategy to another means that the first strategy can replace the second. Under the undirected rule (left), every strategy can replace every other strategy. Under the clockwise rule (middle), each strategy can replace only the strategy it beats in the game: Rock replaces Scissors, Scissors replaces Paper, and Paper replaces Rock. Under the counterclockwise rule (right), each strategy can replace only the strategy that beats it: Rock replaces Paper, Paper replaces Scissors, and Scissors replaces Rock.

Let *f_X_* denote the frequency of strategy *X* (*X* ∈ {R, P, S}) in the population. The expected payoffs of Rock, Paper, and Scissors are

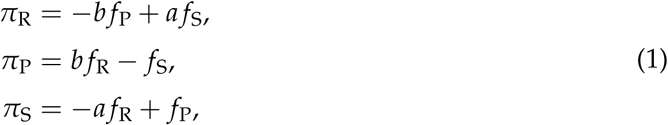

and the mean payoff in the population is 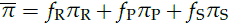.

Strategies spread by imitation. In each time step, an individual *i* is chosen at random to update its strategy, *s_i_*, and another individual *j* is chosen at random as a role model, following the pairwise comparison updating rule (Szabó & To″ke 1998; Traulsen et al. 2006, 2007). Under the classical, undirected rule, *i* adopts the strategy of *j* with a probability that increases with the payoff difference *π_sj_* − *π_si_*. We add a constraint on which strategy can replace which (Fig. 1b). The constraint is described by a 3 × 3 replacement matrix *A* = (*a_XY_*)*_X_*_,*Y*∈{_*_R_*_,*P*,*S*}_, in which *a_XY_* = 1 if strategy *X* can replace strategy *Y*, and *a_XY_* = 0 otherwise. The probability that *i* adopts the strategy of *j* is then

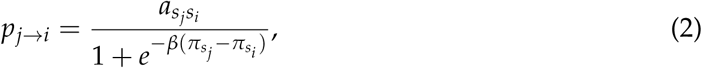

where the intensity of selection, *β* ≥ 0, sets how strongly payoff differences bias imitation. When *a_XY_* = 1 for all *X* and *Y*, every strategy can replace every other, and the model reduces to classical undirected updating (Fig. 1b, left). We compare undirected updating with two directed rules, in which each strategy can replace only one of the other two strategies. Under the clockwise rule, each strategy can replace only the strategy it beats in the game: Rock replaces Scissors, Scissors replaces Paper, and Paper replaces Rock, and none of the reverse replacements occurs (Fig. 1b, middle). Under the counterclockwise rule, each strategy can replace only the strategy that beats it: Rock replaces Paper, Paper replaces Scissors, and Scissors replaces Rock (Fig. 1b, right).

In the limit of a large population, with time measured in units of *N* updates, the frequency of strategy *X* changes according to

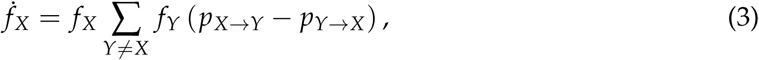

where *p_X_*_→*Y*_ is the probability that an individual using strategy *Y* adopts strategy *X* from a role model using *X* (Eq. 2). We derive Eq. 3 in SI Section 1. In the main text we study weak selection, *β* ≪ 1: we expand the imitation probability in Eq. 2 to first order in *β* to obtain analytical results, and we compute numerical results from the full equations (Eq. 3) at *β* = 0.01. SI Section 2 reports results under strong selection, *β* = 1.

Under undirected updating, the first-order expansion yields the classical replicator equation,

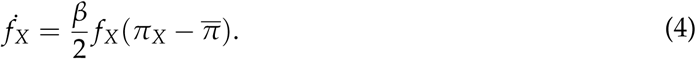

Under clockwise updating, *a*_SR_ = *a*_PS_ = *a*_RP_ = 0, and the replicator equations become

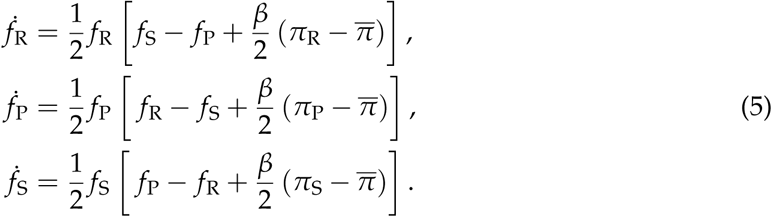

Under counterclockwise updating, *a*_RS_ = *a*_SP_ = *a*_PR_ = 0, and the replicator equations become

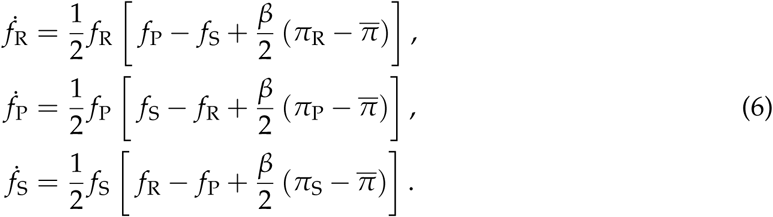

Directed updating changes the replicator equations in two ways (Eqs. 4–6). Under undirected updating, the rate of change of the frequency of strategy *X* depends only on its frequency, *f_X_*, and on its payoff relative to the population mean, 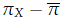. Under directed updating, the rate of change of *f_X_* also depends on a frequency difference between the other two strategies: the frequency of the strategy that *X* can replace, minus the frequency of the strategy that can replace *X*. This frequency-difference term does not depend on the intensity of selection, *β*. In addition, the payoff term has coefficient *β*/4 under directed updating, compared with *β*/2 under undirected updating. In other words, under directed updating the frequency of a strategy changes in response not only to its own payoff but also to the frequencies of the strategy it can replace and of the strategy that can replace it.

## 3 Results

### 3.1 Under directed updating, all three strategies remain at intermediate frequencies

We first compare the trajectories of strategy frequencies under undirected and directed updating, starting from equal frequencies of the three strategies (Fig. 2). When the payoff of Paper against Rock equals the other two payoffs, *a* = *b* = 1, the replicator equations keep the frequencies at 1/3 under all three rules (Fig. 2a). Under undirected updating, as the payoff of Paper against Rock, *b*, increases, the frequency of Scissors remains close to one for most of each oscillation, and the frequencies of Rock and Paper rise only during brief, periodic excursions (Fig. 2a, top row). Under both directed rules, by contrast, the frequencies of all three strategies oscillate periodically, and they remain between 0.05 and 0.6 for every value of *b* shown (Fig. 2a, middle and bottom rows). Compared with undirected updating, the oscillations under directed updating have a shorter period and a smaller amplitude. Because the frequency-difference term does not depend on *β*, the frequencies change on a shorter time scale under directed updating than under undirected updating, in which every term is proportional to *β*. In Monte Carlo simulations of a population of *N* = 10^5^ individuals, the directed rules produce the same oscillations as the replicator equations (Fig. 2b). Under undirected updating, by contrast, a single strategy comes close to fixation in the simulations with large *b*: Scissors when *b* = 20 and *b* = 100, with the other two strategies remaining at low frequencies (Fig. 2b, top row). At *b* = 1, all three strategies remain present over the displayed interval (Fig. 2b, top left).

**Figure 2:**
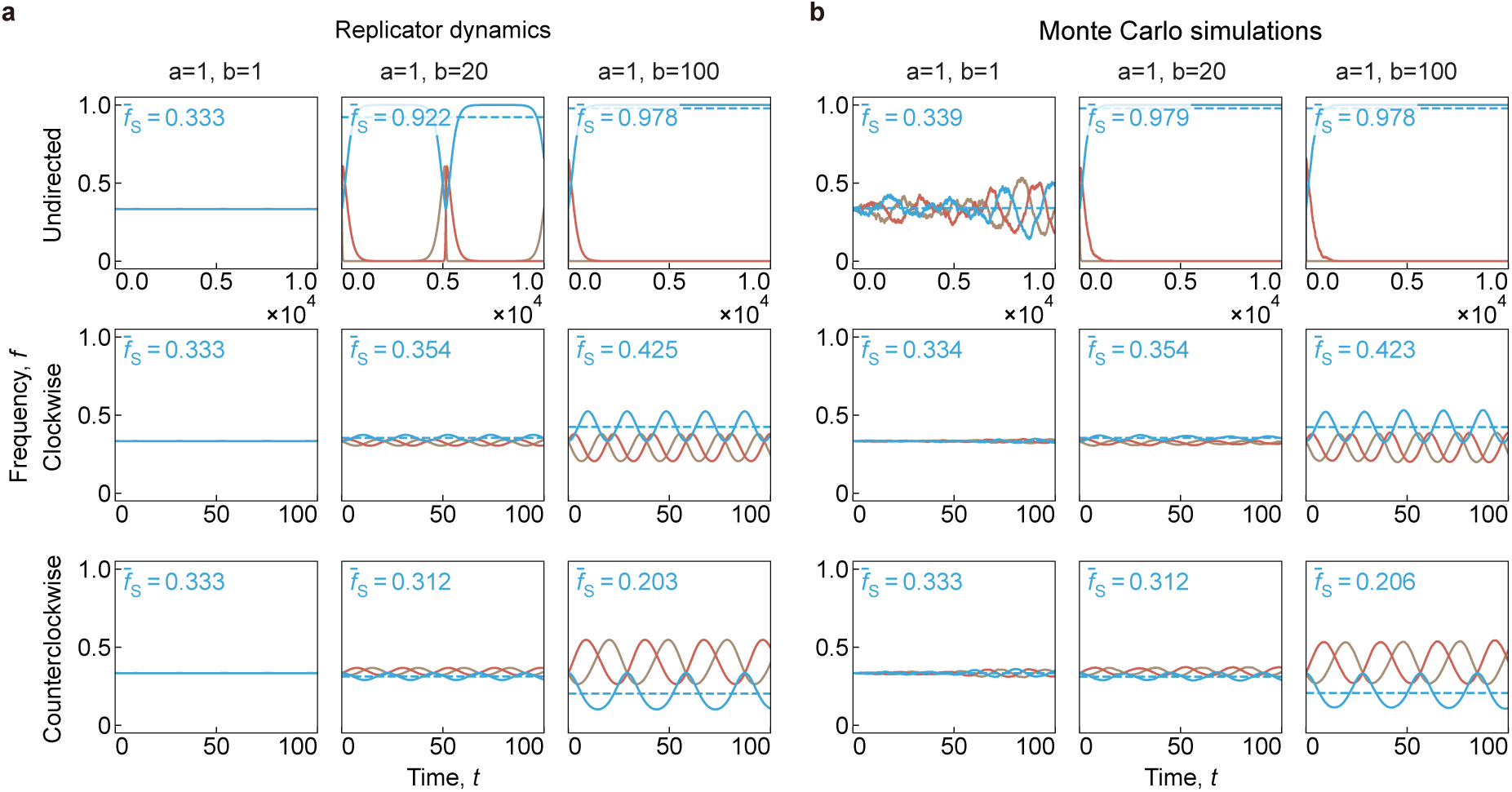
Under directed updating, all three strategies remain at intermediate frequencies. Frequencies of Rock (R, brown), Paper (P, red), and Scissors (S, blue) over time, starting from equal frequencies, under undirected (top row), clockwise (middle row), and counterclockwise (bottom row) updating, for three values of the payoff of Paper against Rock, *b* (columns). **a**, Numerical solutions of the replicator equations (Eqs. 4–6). **b**, Monte Carlo simulations in a finite population. Dashed lines and the values *f*_S_ show the mean frequency of Scissors over the time shown. Under undirected updating, when *b* is large, the frequency of Scissors remains close to one for most of the time in the replicator equations, and Scissors comes close to fixation in the simulations. Under both directed rules, the frequencies of the three strategies oscillate around an interior state. The mean frequency of Scissors increases with *b* under clockwise updating, as it does under undirected updating, and it decreases with *b* under counterclockwise updating. The time axis spans 10^4^ time units for undirected updating and 100 time units for directed updating. Parameters: *a* = 1, *β* = 0.01, and population size *N* = 10^5^ in the simulations.

One might expect that restricting the direction of replacement would make the fixation of one strategy more likely. Instead, the strategy frequencies under directed updating remain closer to their initial values than under undirected updating. We attribute the smaller amplitude of oscillation to the two differences between the replicator equations. First, the payoff term has a smaller coefficient under directed updating (*β*/4, Eqs. 5 and 6) than under undirected updating (*β*/2, Eq. 4), so that payoff differences have a smaller effect on the change in frequencies. Second, and we expect more importantly, directed updating adds the frequency-difference term to the equation for each strategy.

There is a simple way to see how the frequency-difference term produces oscillations, under clockwise updating with *b >* 1 and *a* = 1 (Fig. 2, middle row). At equal frequencies, Paper has the highest payoff and Rock the lowest, so the frequency of Paper increases and the frequency of Rock decreases. Under the clockwise rule, Scissors replaces Paper and is replaced by Rock, and so the frequency-difference term in the equation for Scissors, *f*_P_ − *f*_R_, becomes positive and the frequency of Scissors, *f*_S_, increases (Eq. 5). As *f*_S_ increases, the frequency of Rock, which replaces Scissors, also increases, and *f*_P_ − *f*_R_ decreases. The frequency of Scissors reaches its maximum when *f*_R_ − *f*_P_ equals the payoff term *β*(*π*_S_ − *π*)/2, and it declines thereafter. The same sequence then occurs for Rock and for Paper in turn. And so the frequencies of the three strategies rise and fall in sequence, and they oscillate around an interior state.

### 3.2 Counterclockwise updating reverses the effect of the payoff *b* on the equilibrium

Directed updating also changes how the mean frequency of each strategy depends on the payoff of Paper against Rock, *b* (Fig. 2). Under undirected and clockwise updating, the mean frequency of Scissors increases with *b*. Under counterclockwise updating, the mean frequency of Scissors decreases as *b* increases (Fig. 2, bottom row). The increase under undirected updating is the classical behavior of a Rock-Paper-Scissor game. The interior equilibrium of the undirected replicator equation (Eq. 4) is the Nash equilibrium of the game, 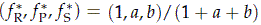, at which the frequency of Scissors increases with *b* even though *b* is a payoff to Paper. Counterclockwise updating reverses this dependence.

The vector fields of the replicator equations show a corresponding difference in the direction of rotation (Fig. 3a). Under undirected updating, the population state rotates around the interior equilibrium in the order of the cycle of dominance in the game: from a population of mostly Scissors to mostly Rock, then to mostly Paper, and back to mostly Scissors (Fig. 1a). Clockwise updating preserves this order of rotation. Counterclockwise updating reverses it, because under this rule each strategy can replace only the strategy that beats it in the game. In other words, under counterclockwise updating the direction of rotation is set by the updating rule, and it is opposite to the cycle of dominance in the payoffs.

**Figure 3:**
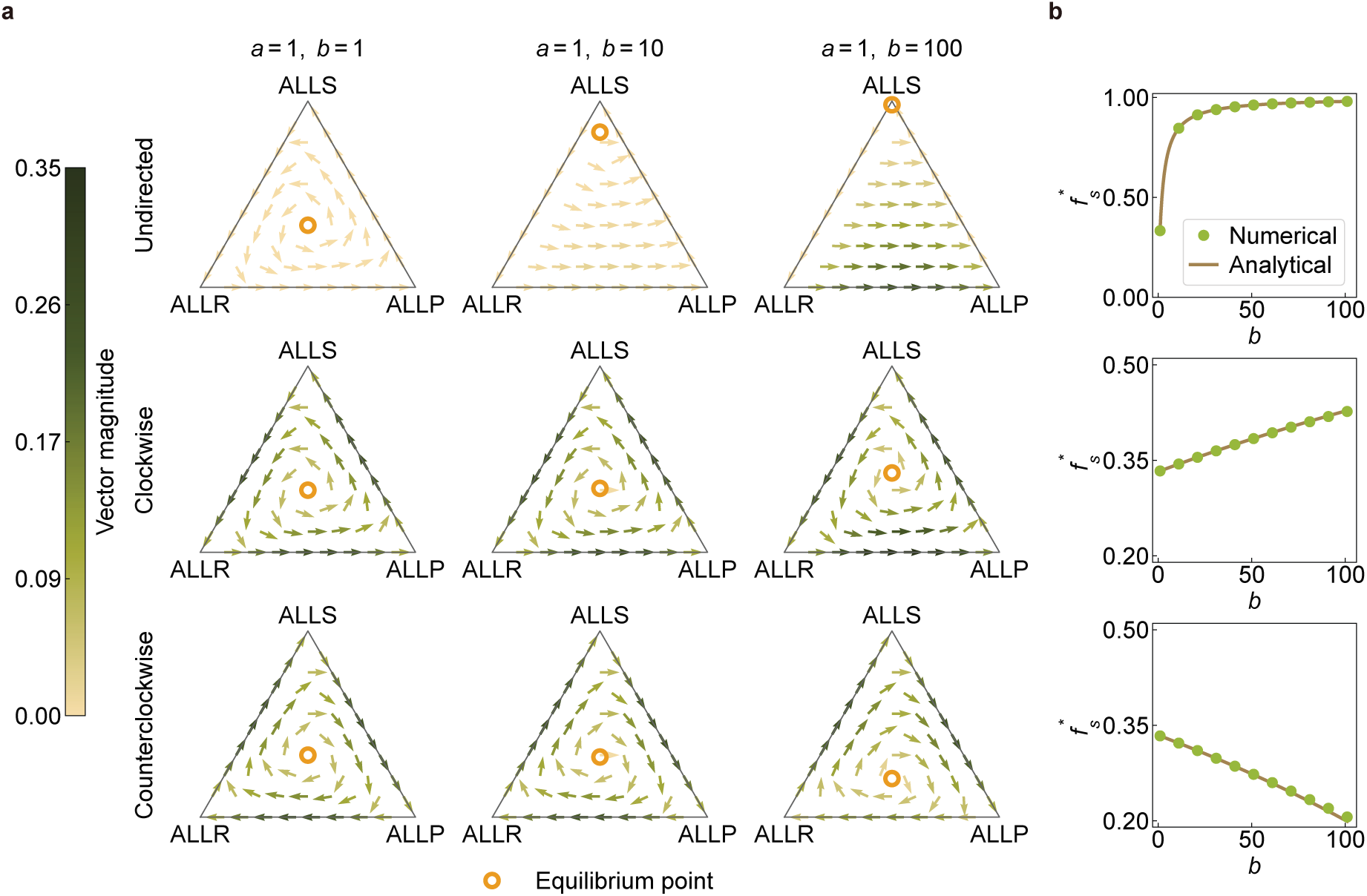
Counterclockwise updating reverses the direction of rotation and the dependence of the interior equilibrium on the payoff *b*. **a**, Vector fields of the full Fermi equations (Eq. 3) on the simplex of strategy frequencies, under undirected (top), clockwise (middle), and counterclockwise (bottom) updating, for three values of the payoff of Paper against Rock, *b* (columns). Each vertex corresponds to a population in which every individual uses the same strategy (ALLR, ALLP, ALLS). Arrows show the direction of change, and their color shows its speed. Open circles mark the interior equilibria. Under undirected and clockwise updating, the population state rotates around the interior equilibrium in the same direction; under counterclockwise updating, it rotates in the opposite direction. **b**, Equilibrium frequency of Scissors, 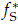, as a function of *b*. Green circles show equilibria computed numerically from the full Fermi equations (Eq. 3), and lines show analytical predictions from the first-order weak-selection approximation (SI Eq. S16). Under undirected and clockwise updating, 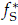 increases with *b*, as it does at the Nash equilibrium of the game; under counterclockwise updating, 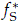 decreases with *b*. The vertical scale of the top panel differs from that of the other two. Parameters: *a* = 1, *β* = 0.01.

The interior equilibrium of the counterclockwise replicator equations shows the reversed dependence on *b* (Fig. 3b). The interior equilibria of Eqs. 4–6 have closed forms (SI Eq. S16). Under undirected and clockwise updating, the equilibrium frequency of Scissors, 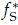, increases with *b*, as it does at the Nash equilibrium of the game. Under counterclockwise updating, 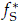 decreases with *b* whenever *β*(1 + *a*) *<* 4, which includes weak selection. And so, under counterclockwise updating, a larger payoff of Paper against Rock lowers the equilibrium frequency of Scissors, which is the opposite of its effect at the Nash equilibrium.

### 3.3 Replacement rates at a reference state explain why the equilibrium frequency of Scissors increases with *b* under undirected and clockwise updating

The dependence of the interior equilibrium on the payoff of Paper against Rock, *b*, can be understood by examining the replacement rates between strategies at a fixed population state. Throughout this section, we set *a* = 1. We consider the reference state **x**_0_ = ( *f*_R_, *f*_P_, *f*_S_) = (*x*_0_, *x*_0_, 1 − 2*x*_0_), with *x*_0_ *<* 1/3, in which Rock and Paper are equally frequent and Scissors is more frequent than either (Fig. 4a). The reference state **x**_0_ is an equilibrium for a given value of *b* when all three strategy frequencies have zero rate of change.

**Figure 4:**
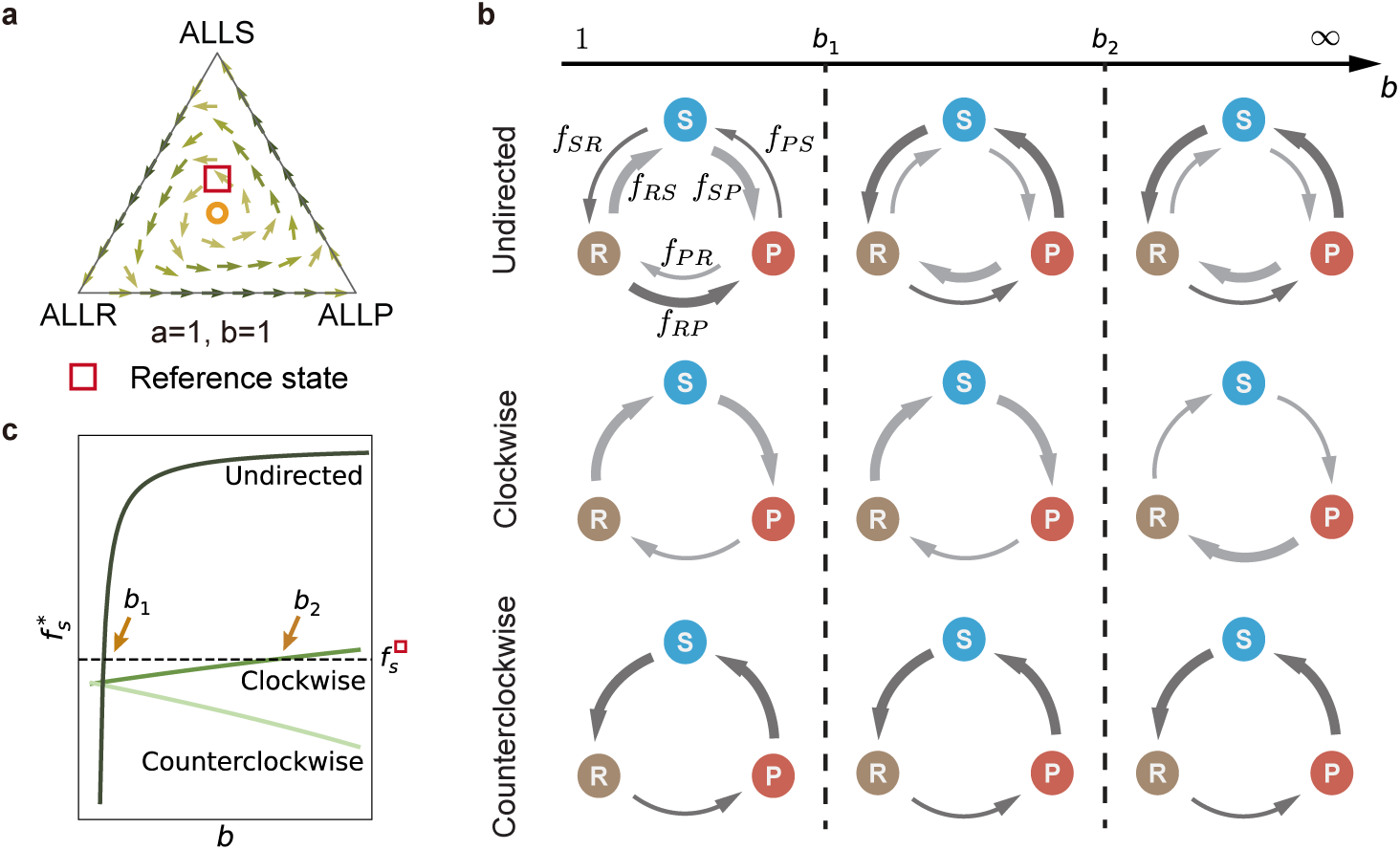
Replacement rates at a reference state explain why the equilibrium frequency of Scissors increases with the payoff *b* under undirected and clockwise updating. **a**, The reference state **x**_0_ (square) and the interior equilibrium (circle) on the vector field of the replicator equations under clockwise updating, for *a* = *b* = 1. **b**, Replacement rates between each pair of strategies at the reference state **x**_0_ = (*x*_0_, *x*_0_, 1 − 2*x*_0_), under undirected (top), clockwise (middle), and counterclockwise (bottom) updating, in three ranges of the payoff of Paper against Rock, *b*, separated by the thresholds *b*_1_ and *b*_2_ (vertical dashed lines). An arrow from one strategy to another means that the first strategy replaces the second, as in Fig. 1b, and the thickness of an arrow indicates the relative size of the replacement rate. Under undirected updating, the sign of the rate of change of the frequency of Rock at the reference state reverses at *b* = *b*_1_, and under clockwise updating it reverses at *b* = *b*_2_. Under counterclockwise updating with weak selection, it does not reverse. **c**, Equilibrium frequency of Scissors, 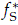, as a function of *b* under the three updating rules. The dotted line marks the frequency of Scissors at the reference state, 1 − 2*x*_0_. Arrows mark the values of *b* at which the reference state is the interior equilibrium: *b*_1_ under undirected updating and *b*_2_ under clockwise updating. Under counterclockwise updating with weak selection, the equilibrium frequency of Scissors does not reach 1 − 2*x*_0_.

At the reference state **x**_0_, the payoffs of the three strategies are

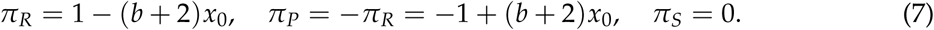

Let *f_XY_* denote the rate at which *X* replaces *Y* at **x**_0_. Note that a replacement rate *f_XY_* carries two subscripts, whereas a frequency *f_X_* carries one. This rate is zero if *X* cannot replace *Y*. For allowed replacements, the rates are

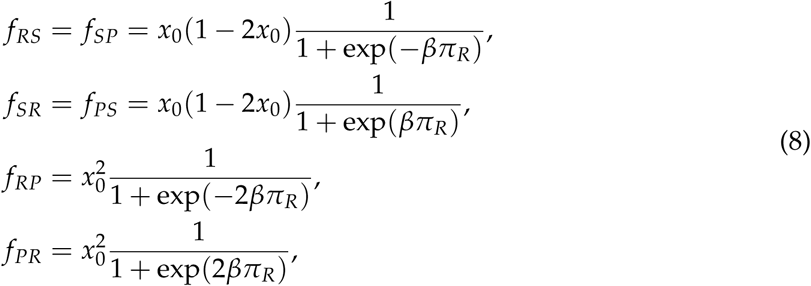

where the subscripts R, P, and S refer to Rock, Paper, and Scissors.

Under undirected updating, all six replacement rates are nonzero. For every value of *b*, the frequency of Scissors does not change at **x**_0_, because *f_RS_* = *f_SP_* and *f_SR_* = *f_PS_*. The replacement rates between Rock and Paper depend on *b*. When *b* = 1, Rock has a positive payoff at the reference state, *π_R_ >* 0, so *f_RP_ > f_PR_*, *f_RS_ > f_SR_*, and *f_SP_ > f_PS_*, and so the frequency of Rock increases and the frequency of Paper decreases. Along the edge of the simplex on which Scissors is absent, by contrast, the frequency of Paper increases and the frequency of Rock decreases. Because the population state rotates around the interior equilibrium, an increase in the frequency of Rock and a decrease in the frequency of Paper at **x**_0_ indicate that the reference state has a higher frequency of Scissors than the interior equilibrium.

Under undirected updating, as the payoff of Paper against Rock increases to *b*_1_ = 1/*x*_0_ − 2, which exceeds 1, the payoff of Rock at the reference state decreases to *π_R_* = 0. Then *f_PR_* = *f_RP_*, *f_SR_* = *f_RS_*, and *f_PS_* = *f_SP_*, the rate of change of the frequency of each strategy is zero, and the reference state **x**_0_ is the interior equilibrium. For *b > b*_1_, *f_PR_ > f_RP_*, *f_SR_ > f_RS_*, and *f_PS_ > f_SP_*, and so at **x**_0_ the frequency of Paper increases and the frequency of Rock decreases, and the reference state has a lower frequency of Scissors than the interior equilibrium. Hence, under undirected updating, the equilibrium frequency of Scissors increases with *b*.

Under clockwise updating, each strategy can replace only the strategy it beats, and so the only nonzero replacement rates at the reference state are *f_RS_*, *f_SP_*, and *f_PR_*. The rate of change of the frequency of Scissors is again zero for every value of *b*, because *f_RS_* = *f_SP_*. When *b* = 1 and selection is weak, *f_PR_< f_RS_*, so that at **x**_0_ the frequency of Rock increases and the frequency of Paper decreases, and the reference state has a higher frequency of Scissors than the interior equilibrium. As *b* increases, *f_PR_* increases and *f_RS_* decreases, and the two rates are equal at a threshold *b*_2_, which exceeds *b*_1_. At *b* = *b*_2_ the rate of change of the frequency of each strategy is zero, and the reference state is the interior equilibrium. For *b > b*_2_, at **x**_0_ the frequency of Paper increases and the frequency of Rock decreases, and the reference state has a lower frequency of Scissors than the interior equilibrium. And so the equilibrium frequency of Scissors under clockwise updating also increases with *b*, but it is lower than under undirected updating for *b >* 1, because the threshold *b*_2_ at which a given reference state is the equilibrium exceeds *b*_1_.

Under counterclockwise updating, each strategy can replace only the strategy that beats it, and so the only nonzero replacement rates are *f_SR_*, *f_PS_*, and *f_RP_*. The rate of change of the frequency of Scissors is zero for every value of *b*, because *f_SR_* = *f_PS_*. When *b* = 1 and selection is weak, *f_RP_ < f_SR_*, so that at **x**_0_ the frequency of Paper increases and the frequency of Rock decreases. As *b* increases, *f_RP_* decreases and *f_SR_* increases, so that *f_RP_ < f_SR_* for every *b >* 1, and the sign of the rate of change of the frequency of Rock at the reference state never changes. In other words, under counterclockwise updating with weak selection, no reference state with *x*_0_ *<* 1/3 becomes the interior equilibrium as *b* increases from 1. The argument that produces an increasing equilibrium frequency of Scissors under undirected and clockwise updating therefore does not apply. The closed-form interior equilibrium of Eq. 6 (SI Eq. S16) shows that the equilibrium frequency of Scissors instead decreases with the payoff of Paper against Rock, *b*, whenever *β*(1 + *a*) *<* 4 (Fig. 3b).

The decrease in the equilibrium frequency of Scissors with *b* under counterclockwise updating also occurs under strong selection, *β* = 1 (Fig. S2). For sufficiently strong selection, however, the inequality *f_RP_ < f_SR_* at *b* = 1 can reverse, and the argument above no longer applies.

## 4 Discussion

We have studied how a restriction on the direction of strategy replacement changes evolutionary dynamics in a generalized Rock-Paper-Scissor game. Under undirected updating, increasing the payoff of Paper against Rock produces a population in which Scissors is close to fixation. Under both directed rules, in which each strategy can replace only one of the other two, the frequencies of all three strategies oscillate around an interior state and remain at intermediate values. And when each strategy can replace only the strategy that beats it, a larger payoff of Paper against Rock lowers the equilibrium frequency of Scissors, which is the opposite of its effect at the Nash equilibrium of the game. This reversal also occurs under strong selection, *β* = 1.

In classical replicator dynamics, the interior equilibrium is the Nash equilibrium, and so it is determined by the payoff matrix alone. Under directed updating, the interior equilibrium depends on the replacement matrix as well. This dependence arises because directed updating adds a term to the replicator equation for each strategy, the frequency of the strategy it can replace minus the frequency of the strategy that can replace it, and this term does not depend on the intensity of selection. Under weak selection, the order in which the strategies rise and fall is therefore set by the updating rule. When the updating rule and the payoffs specify the same cyclic order, as under the clockwise rule, the equilibrium frequency of Scissors increases with the payoff of Paper against Rock, as at the Nash equilibrium. When they specify opposite orders, as under the counterclockwise rule, the population state rotates in the order set by the updating rule, and the equilibrium frequency of Scissors decreases with that payoff. In other words, when replacement is restricted in direction, the payoff matrix alone may not predict even the sign of the change in an equilibrium frequency when a payoff changes.

Directional restrictions on replacement arise in many biological and social systems. In a food chain, a predator can replace its prey but the prey cannot replace the predator (Volterra 1926). In cultural transmission, behaviors are often copied from individuals of higher status or greater experience, and seldom in the reverse direction (Henrich & Gil-White 2001). The clockwise rule describes systems in which a type can displace only the type it outcompetes, as in cyclic competition among microbial strains (Kerr et al. 2002; Kirkup & Riley 2004; Nahum et al. 2011). The counterclockwise rule, in which each strategy can replace only the strategy that beats it in the game, describes a restriction on replacement whose direction is opposite to the direction of payoff dominance. We speculate that such restrictions occur when the ability of one type to replace another depends on traits other than those that determine payoffs in the game, such as the direction in which information flows among individuals.

Prior work has studied restrictions on who can replace whom by placing individuals on a graph, including directed graphs, on which one individual can replace another but not the reverse (Lieberman et al. 2005; Masuda 2009; Masuda & Ohtsuki 2009). In those models the restriction depends on the positions of individuals, and every strategy can in principle replace every other. In our model, by contrast, the population is well-mixed, and the restriction depends on the strategies themselves. Studies of cyclic dominance in Rock-Paper-Scissor games have analyzed how payoffs, mutation, and spatial structure affect the coexistence of strategies (Frean & Abraham 2001; Reichenbach et al. 2007; Mobilia 2010; Szolnoki et al. 2014; Toupo & Strogatz 2015); under undirected updating, our model reduces to the classical replicator equation for such games (Eq. 4).

Several assumptions restrict the scope of our results on directed updating. Our model is deliberately simple: a well-mixed population, a single generalized Rock-Paper-Scissor game, and two replacement matrices in which each strategy can replace exactly one other. The purpose of a model this simple is not to describe any particular ecological or social system, but to identify how the direction of replacement, separately from the payoffs, governs which strategies are common. We analyzed weak selection in the main text and checked the reversal under counterclockwise updating only at *β* = 1. The argument based on replacement rates at a reference state explains why the equilibrium frequency of Scissors increases with *b* under undirected and clockwise updating, but for counterclockwise updating it shows only that this explanation does not apply, and it does not by itself establish the decrease. The decrease follows instead from the closed-form equilibrium of the first-order equations (SI Eq. S16). Likewise, our account of the smaller amplitude of oscillation under directed updating is based on a comparison of the terms in the replicator equations, not on a derivation. We expect that replacement matrices in which only some reverse replacements are forbidden would produce behavior intermediate between undirected updating and the two directed rules, but we have not analyzed them.

Theories of strategy evolution typically predict equilibrium frequencies from the payoff matrix of the game. Our results suggest that, wherever the direction of replacement is constrained, a full account of evolutionary outcomes requires the replacement matrix as well as the payoffs.

## Supplementary Information

Section 1 derives the replicator equations of the main text for an arbitrary replacement matrix and expands them under weak selection. Section 2 reports numerical solutions and simulations under strong selection, *β* = 1.

## 1 Replicator dynamics

In this section we derive the replicator equations of the main text (Eqs. 3–6) for pairwise comparison updating with an arbitrary replacement matrix *A* = (*a_XY_*), in which *a_XY_* = 1 if strategy *X* can replace strategy *Y* and *a_XY_*= 0 otherwise. We first consider a finite population of *N* individuals, and we let *f_X_* denote the frequency of strategy *X* ∈ {R, P, S} in the population. Each update involves two individuals chosen at random: a *focal* individual, who may change its strategy, and a *role model*, whose strategy the focal individual may adopt. In each update, the frequency of strategy *X* changes through one of two events:

- **Increase**: An individual using strategy *Y* = *X* is chosen to update its strategy, with probability *f_Y_*. This individual chooses a role model using strategy *X*, with probability *f_X_*, and adopts strategy *X* with probability *p_X_*_→*Y*_, which increases *f_X_* by 1/*N*.
- **Decrease**: An individual using strategy *X* is chosen to update its strategy, with probability *f_X_*. This individual chooses a role model using strategy *Y* = *X*, with probability *f_Y_*, and adopts strategy *Y* with probability *p_Y_*_→*X*_, which decreases *f_X_* by 1/*N*.

The expected change in the frequency of strategy *X* in a single update is therefore

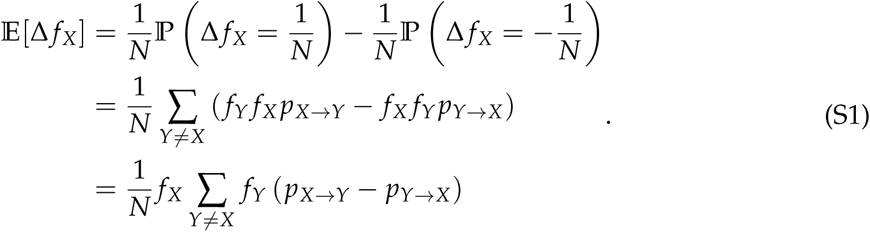

Under pairwise comparison updating with replacement matrix *A*, the probability that an individual using strategy *Y* adopts strategy *X* from a role model using *X* is

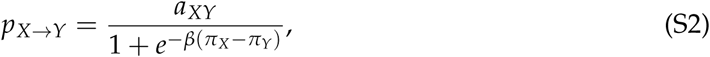

where *π_X_* is the expected payoff of strategy *X* (Eq. 1 of the main text) and *β* ≥ 0 is the intensity of selection. Substituting into Eq. S1 gives

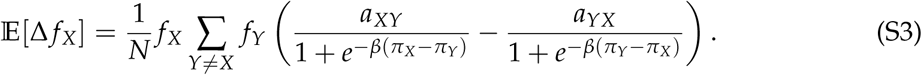

Measuring time in units of *N* updates and taking the limit of a large population, we obtain from Eq. S3 the replicator equation for an arbitrary replacement matrix and an arbitrary intensity of selection,

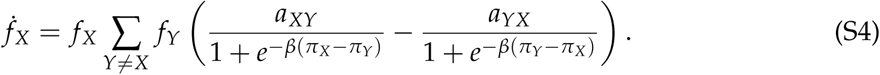

### 1.1 Undirected updating rule

Under undirected updating, every strategy can replace every other strategy, that is, *a_XY_* = 1 for all strategies *X* and *Y*, and Eq. S4 becomes

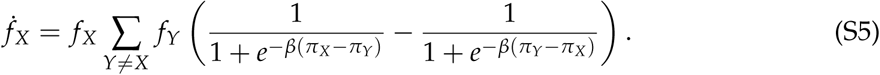

Under weak selection, *β* ≪ 1, the imitation probability has the expansion

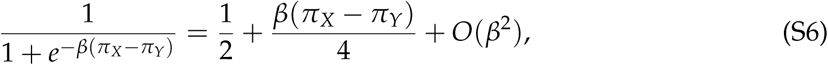

and substituting it, with the mean payoff *π* = *f_R_π_R_* + *f_P_π_P_* + *f_S_π_S_*, gives

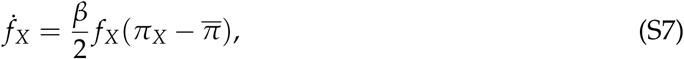

which is the classical replicator equation of evolutionary game theory (Eq. 4 of the main text).

### 1.2 Directed updating rules

We consider two directed updating rules, the clockwise rule and the counterclockwise rule. Under the clockwise rule, each strategy can replace only the strategy it beats in the game: Rock replaces Scissors, Scissors replaces Paper, and Paper replaces Rock, and none of the reverse replacements occurs. The replacement matrix is

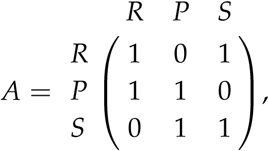

and Eq. S4 becomes, for arbitrary intensity of selection *β*,

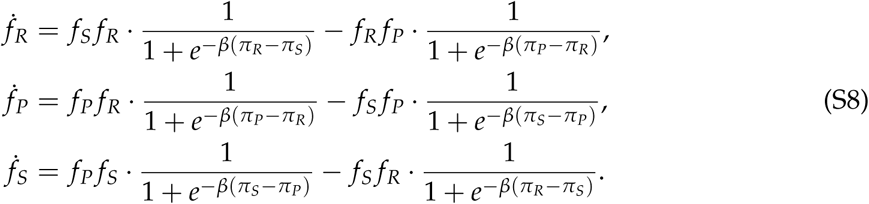

Under weak selection, *β* ≪ 1, the two imitation probabilities in the equation for *f_R_* have the expansions

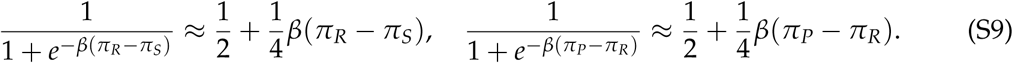

Substituting into *f_R_* gives

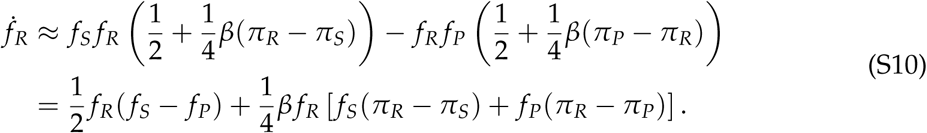

Because *f_S_*+ *f_P_* = 1 − *f_R_*, the term in square brackets equals

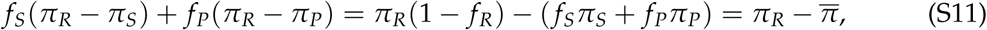

where the second equality uses the definition of the mean payoff, *π* = *f_R_π_R_* + *f_P_π_P_* + *f_S_π_S_*. Hence Eq. S10 becomes

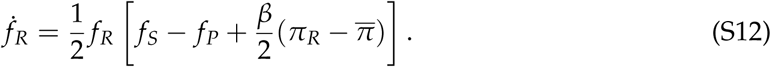

The equations for Paper and Scissors follow in the same way:

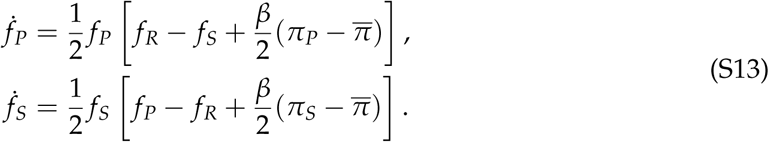

Under the counterclockwise rule, each strategy can replace only the strategy that beats it in the game: Rock replaces Paper, Paper replaces Scissors, and Scissors replaces Rock, and none of the reverse replacements occurs. The replacement matrix is

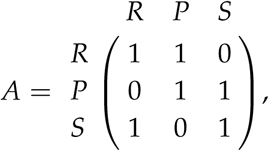

and Eq. S4 becomes, for arbitrary intensity of selection *β*,

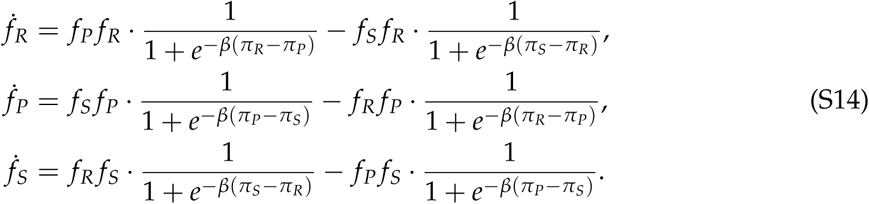

Under weak selection, the same expansion as for the clockwise rule gives

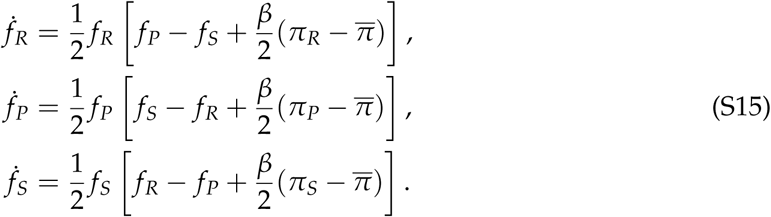

### 1.3 Interior equilibria under weak selection

We derive the interior equilibria of the first-order weak-selection equations used in the main text. Because the game is zero-sum, the mean payoff is *π* = 0 for every state of the population. At an interior equilibrium every frequency is positive, so the term in square brackets in each weak-selection equation equals zero, and each such term is linear in the frequencies, because the payoffs (Eq. 1) are linear. Under clockwise updating, for example, the condition for Rock is 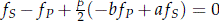. Solving these linear equations together with *f_R_* + *f_P_* + *f_S_* = 1, with *β >* 0, gives the interior equilibrium

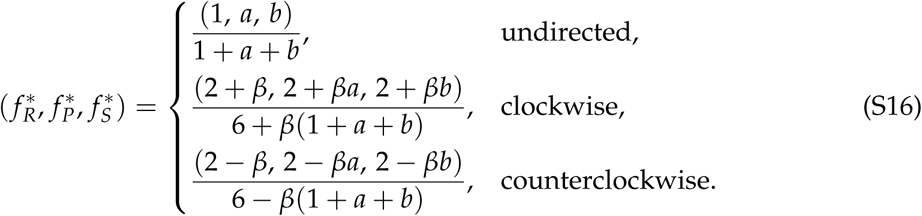

Under counterclockwise updating, the three equilibrium frequencies are positive when *β* max(1, *a*, *b*) *<* 2, which holds under weak selection and throughout Fig. 3b. The lines in Fig. 3b show the equilibrium frequency of Scissors, 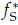, from these expressions.

Differentiating the equilibrium frequency of Scissors with respect to the payoff of Paper against Rock, *b*, gives 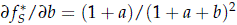 under undirected updating, *β*[4 + *β*(1 + *a*)]/[6 + *β*(1 + *a* + *b*)]^2^ under clockwise updating, and *β*[*β*(1 + *a*) − 4]/[6 − *β*(1 + *a* + *b*)]^2^ under counterclockwise updating. The first two are positive for every *b*. The third is negative whenever *β*(1 + *a*) *<* 4, which includes weak selection.

## 2 Strong selection

The main text analyzes weak selection. Here we set the intensity of selection to *β* = 1 and solve the replicator equation without the weak-selection expansion (Eq. S4), under undirected, clockwise, and counterclockwise updating. We compare these solutions with Monte Carlo simulations in a population of *N* = 10^5^, with *a* = 1 and *b* = 1, 2, 4 (Fig. S1).

Under strong selection, we find the same pattern as under weak selection: the mean frequency of Scissors over the time shown increases with *b* under undirected and clockwise updating, and decreases with *b* under counterclockwise updating. In the replicator equation, the period under counterclockwise updating is longer at *b* = 4 than at *b* = 2, so that a complete cycle is not visible at *b* = 4 within the 100 time units shown. The equilibrium frequency of Scissors shows the same dependence on *b*: it increases under undirected and clockwise updating, and decreases under counterclockwise updating (Fig. S2).

**Figure S1:**
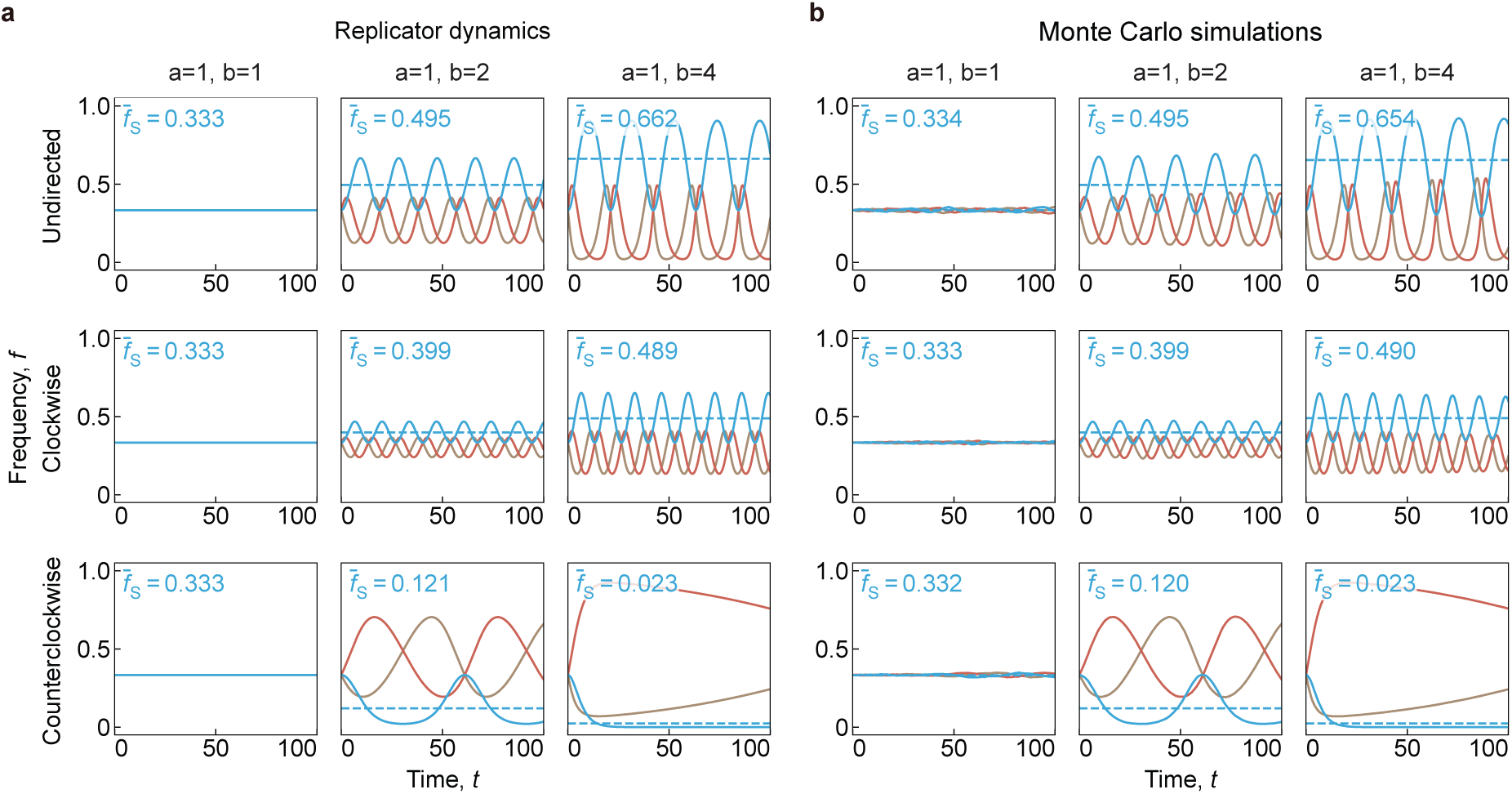
Under strong selection, counterclockwise updating also reverses the dependence of the mean frequency of Scissors on the payoff. Frequencies of Rock (R, brown), Paper (P, red), and Scissors (S, blue) over time, starting from nearly equal frequencies, under undirected (top row), clockwise (middle row), and counter-clockwise (bottom row) updating, for three values of the payoff of Paper against Rock, *b* = 1, 2, 4 (columns). a, Numerical solutions of the replicator equation (Eq. S4). b, Monte Carlo simulations in a finite population, with one realization shown in each panel. Dashed lines and the values *f*_S_ show the mean frequency of Scissors over the time shown. Under strong selection, the mean frequency of Scissors over this interval increases with *b* under undirected and clockwise updating, and decreases with *b* under counterclockwise updating, following the same pattern as under weak selection. The time axis spans 100 time units in all panels. Parameters: *a* = 1, *β* = 1, and population size *N* = 10^5^ in the simulations.

**Figure S2:**
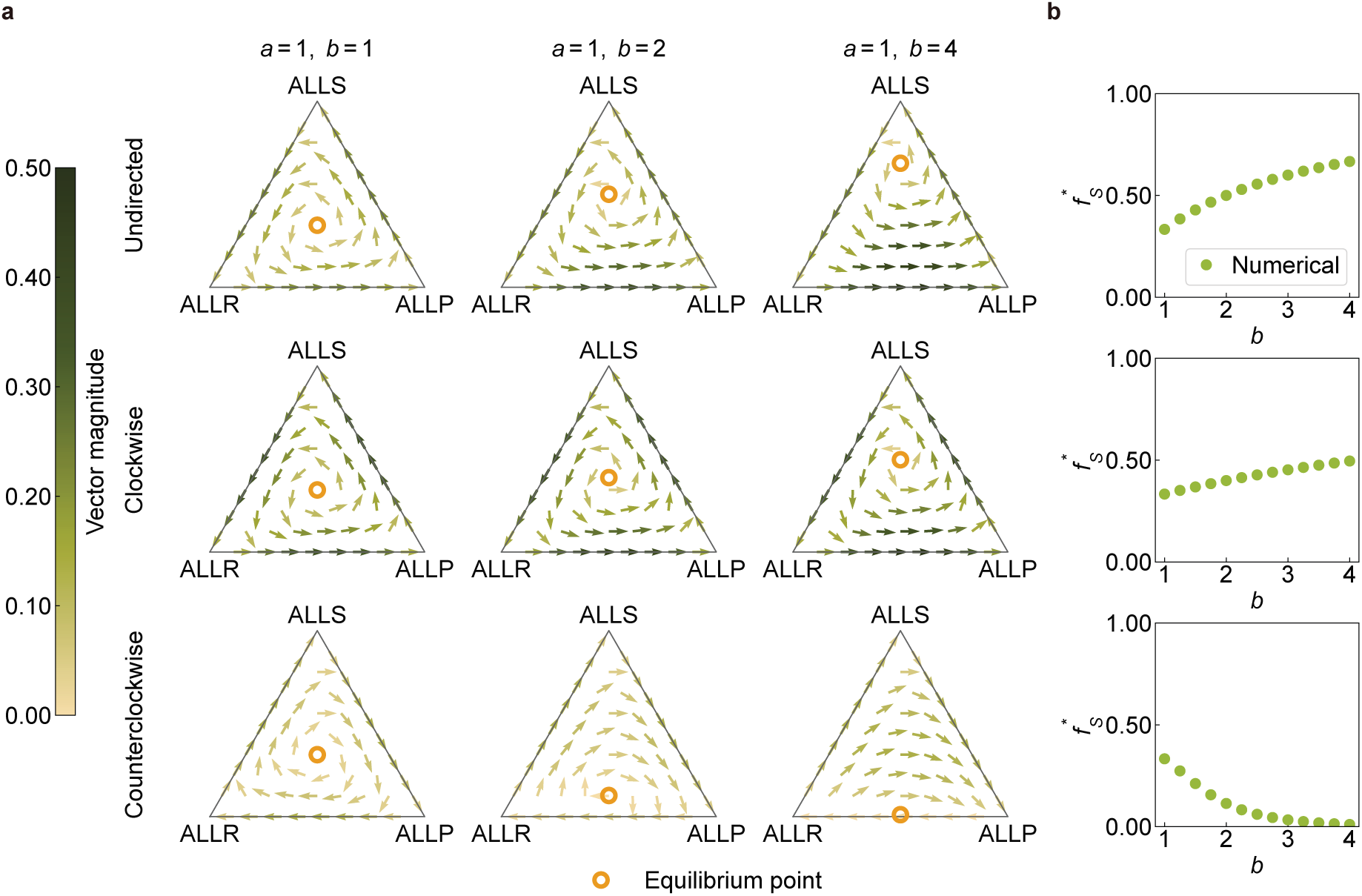
Under strong selection, counterclockwise updating also reverses the dependence of the equilibrium frequency of Scissors on the payoff *b*. **a**, Vector fields of the replicator equation without the weak-selection expansion (Eq. S4) on the simplex of strategy frequencies, under undirected (top), clockwise (middle), and counterclockwise (bottom) updating, with *b* = 1, 2, 4 from left to right. Arrows show the direction of change, and their color shows its speed. Orange open circles mark the interior equilibria. b, Green points are the numerically computed equilibrium frequency of Scissors as a function of *b*, under the corresponding updating rule in each row. This frequency increases with *b* under undirected and clockwise updating, and decreases with *b* under counterclockwise updating. Parameters: *a* = 1, *β* = 1.

## References

1. Allen B, Lippner G, Chen YT, Fotouhi B, Momeni N, et al. (2017). Evolutionary dynamics on any population structure. Nature 544: 227–230. 10.1038/nature21723

2. Anderson RM, May RM (1991). Infectious Diseases of Humans: Dynamics and Control. Oxford: Oxford University Press. 10.1093/oso/9780198545996.001.0001

3. Asano K, Nabeyama A, Miyake Y, Qiu CH, Kurita A, et al. (2011). CD169-positive macrophages dominate antitumor immunity by crosspresenting dead cell-associated antigens. Immunity 34: 85–95. 10.1016/j.immuni.2010.12.011

4. Baghdadi M, Yoneda A, Yamashina T, Nagao H, Komohara Y, et al. (2013). TIM-4 glycoprotein-mediated degradation of dying tumor cells by autophagy leads to reduced antigen presentation and increased immune tolerance. Immunity 39: 1070–1081. 10.1016/j.immuni.2013.09.014

5. Chudek M, Heller S, Birch S, Henrich J (2012). Prestige-biased cultural learning: bystander’s differential attention to potential models influences children’s learning. Evolution and Human Behavior 33: 46–56. 10.1016/j.evolhumbehav.2011.05.005

6. Cressman R, Tao Y (2014). The replicator equation and other game dynamics. Proceedings of the National Academy of Sciences 111: 10810–10817. 10.1073/pnas.1400823111

7. Frean M, Abraham ER (2001). Rock–scissors–paper and the survival of the weakest. Proceedings of the Royal Society B: Biological Sciences 268: 1323–1327. 10.1098/rspb.2001.1670

8. Helbing D (1992). A mathematical model for behavioral changes by pair interactions. In: Haag G, Mueller U, Troitzsch KG (eds.). Economic Evolution and Demographic Change. Lecture Notes in Economics and Mathematical Systems, Volume 395. Berlin, Heidelberg: Springer, pp. 330–348. 10.1007/978-3-642-48808-518

9. Henrich J, Broesch J (2011). On the nature of cultural transmission networks: evidence from Fijian villages for adaptive learning biases. Philosophical Transactions of the Royal Society B: Biological Sciences 366: 1139–1148. 10.1098/rstb.2010.0323

10. Henrich J, Gil-White FJ (2001). The evolution of prestige: Freely conferred deference as a mechanism for enhancing the benefits of cultural transmission. Evolution and Human Behavior 22: 165–196. 10.1016/S1090-5138(00)00071-4

11. Hethcote HW (2000). The mathematics of infectious diseases. SIAM Review 42: 599–653. 10.1137/S0036144500371907

12. Hofbauer J, Sigmund K (1998). Evolutionary Games and Population Dynamics. Cambridge: Cambridge University Press. 10.1017/CBO9781139173179

13. Hofbauer J, Sigmund K (2003). Evolutionary game dynamics. Bulletin of the American Mathematical Society 40: 479–519. 10.1090/S0273-0979-03-00988-1

14. Holling CS (1959). Some characteristics of simple types of predation and parasitism. The Canadian Entomologist 91: 385–398. 10.4039/Ent91385-7

15. Kermack WO, McKendrick AG (1927). A contribution to the mathematical theory of epidemics. Proceedings of the Royal Society of London. Series A 115: 700–721. 10.1098/rspa.1927.0118

16. Kerr B, Riley MA, Feldman MW, Bohannan BJM (2002). Local dispersal promotes biodiversity in a real-life game of rock–paper–scissors. Nature 418: 171–174. 10.1038/nature00823

17. Kirkup BC, Riley MA (2004). Antibiotic-mediated antagonism leads to a bacterial game of rock-paper-scissors in vivo. Nature 428: 412–414. 10.1038/nature02429

18. Lieberman E, Hauert C, Nowak MA (2005). Evolutionary dynamics on graphs. Nature 433: 312–316. 10.1038/nature03204

19. Masuda N (2009). Directionality of contact networks suppresses selection pressure in evolutionary dynamics. Journal of Theoretical Biology 258: 323–334. 10.1016/j.jtbi.2009.01.025

20. Masuda N, Ohtsuki H (2009). Evolutionary dynamics and fixation probabilities in directed networks. New Journal of Physics 11: 033012. 10.1088/1367-2630/11/3/033012

21. Maynard Smith J, Price GR (1973). The logic of animal conflict. Nature 246: 15–18. 10.1038/246015a0

22. Mobilia M (2010). Oscillatory dynamics in rock-paper-scissors games with mutations. Journal of Theoretical Biology 264: 1–10. 10.1016/j.jtbi.2010.01.008

23. Nahum JR, Harding BN, Kerr B (2011). Evolution of restraint in a structured rock-paperscissors community. Proceedings of the National Academy of Sciences 108: 10831–10838. 10.1073/pnas.1100296108

24. Ohtsuki H, Hauert C, Lieberman E, Nowak MA (2006). A simple rule for the evolution of cooperation on graphs and social networks. Nature 441: 502–505. 10.1038/nature04605

25. Reichenbach T, Mobilia M, Frey E (2007). Mobility promotes and jeopardizes biodiversity in rock-paper-scissors games. Nature 448: 1046–1049. 10.1038/nature06095

26. Sandholm WH (2010). Population Games and Evolutionary Dynamics. Cambridge, MA: MIT Press.

27. Schlag KH (1998). Why imitate, and if so, how?: A boundedly rational approach to multiarmed bandits. Journal of Economic Theory 78: 130–156. 10.1006/jeth.1997.2347

28. Schuster P, Sigmund K (1983). Replicator dynamics. Journal of Theoretical Biology 100: 533–538. 10.1016/0022-5193(83)90445-9

29. Sheng A, Su Q, McAvoy A, Wang L, Plotkin JB (2026). Cooperation conflicts with equality when allocating public goods. Nature 655: 1217–1223. 10.1038/s41586-026-10550-3

30. Sinervo B, Lively CM (1996). The rock–paper–scissors game and the evolution of alternative male strategies. Nature 380: 240–243. 10.1038/380240a0

31. Su Q, Allen B, Plotkin JB (2022). Evolution of cooperation with asymmetric social interactions. Proceedings of the National Academy of Sciences 119: e2113468118. 10.1073/pnas.2113468118

32. Szabó G, To″ke C (1998). Evolutionary prisoner’s dilemma game on a square lattice. Physical Review E 58: 69–73. 10.1103/PhysRevE.58.69

33. Szolnoki A, Mobilia M, Jiang LL, Szczesny B, Rucklidge AM, et al. (2014). Cyclic dominance in evolutionary games: a review. Journal of the Royal Society Interface 11: 20140735. 10.1098/rsif.2014.0735

34. Taylor PD, Jonker LB (1978). Evolutionarily stable strategies and game dynamics. Mathematical Biosciences 40: 145–156. 10.1016/0025-5564(78)90077-9

35. Toupo DFP, Strogatz SH (2015). Nonlinear dynamics of the rock-paper-scissors game with mutations. Physical Review E 91: 052907. 10.1103/PhysRevE.91.052907

36. Traulsen A, Claussen JC, Hauert C (2005). Coevolutionary dynamics: From finite to infinite populations. Physical Review Letters 95: 238701. 10.1103/PhysRevLett.95.238701

37. Traulsen A, Nowak MA, Pacheco JM (2006). Stochastic dynamics of invasion and fixation. Physical Review E 74: 011909. 10.1103/PhysRevE.74.011909

38. Traulsen A, Pacheco JM, Nowak MA (2007). Pairwise comparison and selection temperature in evolutionary game dynamics. Journal of Theoretical Biology 246: 522–529. 10.1016/j.jtbi.2007.01.002

39. Traulsen A, Semmann D, Sommerfeld RD, Krambeck HJ, Milinski M (2010). Human strategy updating in evolutionary games. Proceedings of the National Academy of Sciences 107: 2962–2966. 10.1073/pnas.0912515107

40. Volterra V (1926). Fluctuations in the abundance of a species considered mathematically. Nature 118: 558–560. 10.1038/118558a0

41. Zhou Y, Fei M, Zhang G, Liang WC, Lin W, et al. (2020). Blockade of the phagocytic receptor MerTK on tumor-associated macrophages enhances P2X7R-dependent STING activation by tumor-derived cGAMP. Immunity 52: 357–373.e9. 10.1016/j.immuni.2020.01.014

